# Prolactin receptor localization and dynamics: Insights from quantitative imaging and mathematical modeling

**DOI:** 10.1101/2025.10.14.682359

**Authors:** Lynne Cherchia, Scott E. Fraser, Stacey D. Finley, Falk Schneider

## Abstract

Signal transduction through the prolactin receptor (PRLR) is crucial in pancreatic β-cell proliferation, impacting pancreatic homeostasis. PRLR-induced JAK/STAT signaling is dynamic, involving changes in spatial organization of signaling molecules. Thus, the spatial organization of PRLR could have strong implications on signaling output. Internalization has been shown and modeled in other signaling pathways but has not been considered in a mathematical model of PRLR signaling. Here, we use live-cell fluorescence imaging, reconstitution approaches, and fluorescence correlation spectroscopy (FCS) to inform a mathematical model of PRLR signaling. Internal PRLR localization is observed in primary pancreatic tissue and in an engineered PRLR expression system. Our imaging data indicate the presence of intracellular and plasma membrane-bound receptor populations. We use FCS to resolve the membrane-bound PRLR population. Based on our data, we include internalization dynamics within an ordinary differential equation (ODE) model of PRLR signaling. We employ the model to explore how the spatial heterogeneity of PRLR affects downstream signaling. We show that the model is more sensitive to PRLR trafficking rates and ability to promote signaling than to its initial spatial distribution. Our data underscore the versatility of a modeling-imaging framework to quantitatively understand signal transduction in and beyond β-cells.

**Significance Statement:**

- Prolactin receptor (PRLR) signal transduction impacts the growth and survival of insulin-secreting cells, making this pathway a target for building our understanding of pancreatic homeostasis and exploring potential diabetes therapeutics.
- Live fluorescence imaging techniques applied within an engineered PRLR expression platform indicate PRLR localization patterns consistent with primary pancreatic tissue and the presence of two spatially distinct PRLR populations. These observations inform a predictive mathematical model of PRLR signaling.
- Integrating experimental data tailored to computational approaches shapes our understanding of complex, multiscale systems such as signal transduction. A generalizable modeling-imaging framework enables the study of molecular dynamics beyond β-cells.

## Introduction

Pancreatic β-cell mass is a critical factor in diabetes. Both type 1 and type 2 diabetes result from insufficient numbers of functional, insulin-secreting cells, suggesting that a β-cell replacement approach holds promise as a diabetes therapeutic. The objective of restoring functional β-cell mass has led to efforts in pancreatic islet transplantation (Shapiro et al., 2017; Rickels and Robertson, 2019; Cayabyab et al., 2021), derivation of glucose-responsive, β-like cells from human pluripotent stem cells (Pagliuca et al., 2014; Rezania et al., 2014), and α-cell reprogramming (Xiao et al., 2018; Karakose et al., 2024). These approaches are limited by autoimmune attacks (Shapiro et al., 2000; Aguayo-Mazzucato and Bonner-Weir, 2018), and ultimately the scalability of these therapeutics to the millions of individuals currently living with diabetes. Exploring endogenous proliferation pathways addresses these limitations.

β-cell proliferation is known to be controlled in part through the prolactin receptor (PRLR) (Brelje et al., 2002, 2004; Millette et al., 2022). PRLR is a single-pass transmembrane receptor comprising an N-terminal extracellular domain, a transmembrane domain, and a C-terminal intracellular domain. PRLR mediates the effects of lactogenic hormones such as prolactin (PRL) on pancreatic β-cells by inducing Janus kinase 2 and signal transducer and activator of transcription 5 (JAK/STAT) signal transduction. This lactogenic signaling is known to stimulate β-cell replication without compromising the cells’ ability to produce insulin (Friedrichsen et al., 2003). As a result of this signal transduction, there is also evidence of STAT5-dependent expression of Bcl-xL (Fujinaka et al., 2007) and cyclin D2 (Friedrichsen et al., 2003; Sorenson and Brelje, 2009), indicating a direct link between PRLR signal transduction and β-cell proliferation.

To characterize its role in β-cell proliferation, PRLR organization and signaling have been studied in vitro (Friedrichsen et al., 2003), in vivo (Fujinaka et al., 2007; Millette et al., 2022), and in silico (Mortlock et al., 2021). As a transmembrane cytokine receptor, PRLR is expected to localize to the cell surface, following a pattern similar to other single-pass transmembrane proteins such as CD4, CD8, or CD86 (Mørch et al., 2022). However, close inspection of imaging data from the literature suggests intracellular PRLR localization (Tallet et al., 2011; Lopez-Pulido et al., 2013; Araya-Secchi et al., 2023). These prior studies focus on the molecular effects of structural changes to PRLR as well as the cell proliferation effects of PRLR antagonists. By raising questions about PRLR organization, these studies build the foundation for our work probing PRLR organization, and potential trafficking, and hypothesizing about the implications of intracellular PRLR localization. Further, many studies of PRLR signaling employ immunoblotting to detect levels of downstream signaling proteins in fixed tissues (Brelje et al., 2002, 2004; Fujinaka et al., 2007). This approach relies on static timepoints to detect the time-varying levels of signaling proteins. Simulations of in silico models parameterized by these data, in contrast, can predict continuous signaling dynamics over time. Our previously established mathematical model of PRLR-mediated JAK/STAT signaling has been used to explore the effects of feedback modules on STAT5 activation (Mortlock et al., 2021), the role of cell-to-cell heterogeneity in activating a pro-proliferation response following hormone signaling (Simoni et al., 2022), and the application of an improved approach to Bayesian hypothesis formation (Huber et al., 2023). However, this model has not yet been informed by live-cell data and thus has not considered PRLR recycling, storage, or distribution mechanisms implicated by imaging data.

Together, these prior findings motivate the development and application of the next generation of analyses – here, imaging modalities with high spatiotemporal resolution that can be integrated with mathematical modeling to bridge scales – to investigate the effects of intracellular PRLR localization on cell signaling. We apply a combination of high-resolution imaging, spectroscopy, and primary and reconstituted PRLR expression systems in conjunction with mathematical modeling in an effort to make sense of the complex signaling dynamics and their downstream outcomes for β-cell proliferation. Live fluorescence microscopy and spectroscopy offer tools aligned with studying PRLR signaling and capturing signaling dynamics across scales. Live imaging with high temporal resolution provides live measurements of receptor localization, and fluorescence correlation spectroscopy (FCS) methods directly measure molecular characteristics and behavior of signaling proteins. Further, integrating these high-resolution, quantitative imaging modalities with mathematical modeling enables hypothesis testing of time-dependent behaviors in complex systems such as intracellular signaling pathways.

In this work, we take a deep dive into PRLR organization. We develop a reconstituted expression system in which fluorescently labeled PRLR is analyzed with fluorescence imaging and FCS, and PRLR’s observed behavior is then translated into a mathematical representation in our model. Our imaging results point to the importance of a PRLR trafficking mechanism, and our simulations suggest the PRLR signaling model is robust to changes in initial PRLR localization.

## Results

We present our analysis in three primary segments. First, we describe our experimental setup and how we envision its integration with existing and to-be-developed modeling approaches. Next, we present our experimental results on localization and dynamics of endogenous and reconstituted PRLR using fluorescence microscopy and spectroscopy methods. Finally, we present integration of the intracellular pool of PRLR into a revised ODE model of the JAK/STAT signaling pathway.

### Integrating live imaging and ODE modeling

We set out to integrate observations from live imaging experiments into a predictive model of PRLR signaling to better understand the role of subcellular PRLR localization. Figure 1 details the integrated experimental-computational workflow designed to answer this question. The established model of PRLR-mediated JAK/STAT signaling uses ODEs to represent biochemical reactions. PRL binds PRLR-JAK complexes at the cell surface, leading to STAT5 activation in the cytosol and translocation to the nucleus where it acts as a transcription factor promoting expression of target genes and eliciting a response (Figure 1A). Based on imaging data, we propose building on this model by adding an intracellular compartment to account for PRLR recycling and storage (Figure 1B). To more closely study these cycling dynamics, we develop an experimental system consisting of fluorescently labeled PRLR, providing an avenue to measure localization dynamics (Figure 1C). Combining our experimental system and proposed signal transduction mechanism, we analyze PRLR localization and dynamics by applying live labeling and FCS as well as testing PRLR cycling and signal transduction hypotheses within our updated, imaging data-informed mathematical model (Figure 1D).

**Figure 1:**
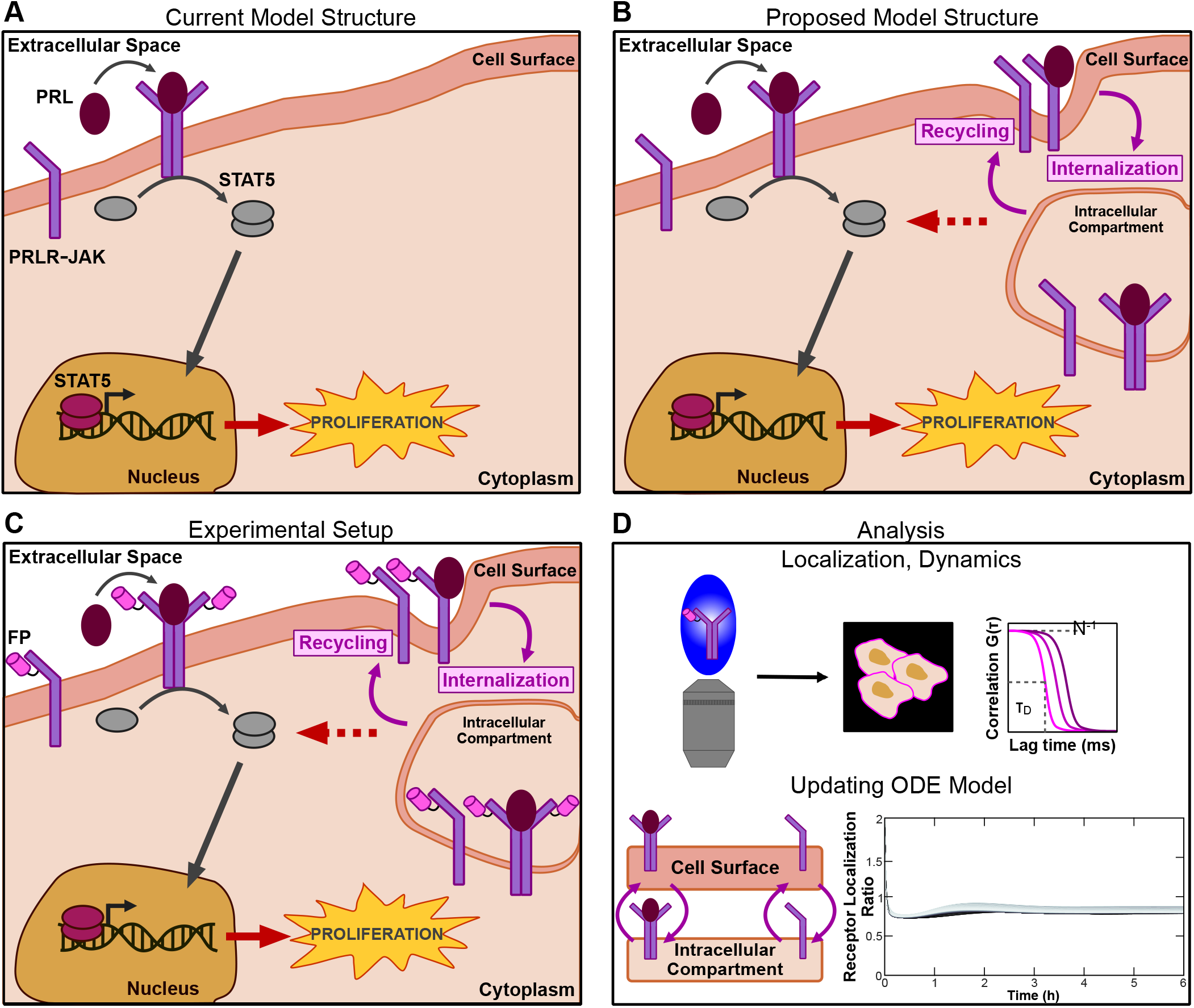
Experimental and computational workflow revises existing mathematical model by integrating imaging and modeling. **A** In PRL-mediated JAK/STAT signaling, PRL binds to PRLR-JAK at the plasma membrane, resulting in receptor dimerization and JAK activation. Activated JAK phosphorylates STAT5 monomers in the cytosol, which then dimerize and translocate to the nucleus. Nuclear STAT5 contributes to the β-cell response to JAK/STAT signaling by acting as a transcription factor promoting the expression of target genes. **B** Proposed model adds recycling and internalization reactions for free and ligand-bound forms of PRLR. **C** The experimental setup fuses PRLR with a fluorescent protein (FP) to allow for visualization and diffusion dynamics studies in a live-cell, reconstituted system of PRLR expression. **D** Quantitative imaging is applied within the reconstituted system to analyze PRLR localization, and these observations inform the structure of the ODE model.

### Immunofluorescence in rodent tissue reveals heterogeneous PRLR localization

To establish a ground truth for PRLR localization in primary tissue, we examine PRLR localization patterns in rodent cells. We perform immunofluorescence against PRLR in INS-1E cells (Figures 2A and 2B) and whole murine islets (Figures 2C and 2D). We use E-cadherin as a counter stain for membrane localization. Confocal fluorescence imaging shows predominantly intracellular PRLR localization in both fixed rat insulinoma cells (Figure 2A) and murine islets (Figure 2C). Fluorescence intensity line profiles are measured across the plasma membrane, in multiple cells at independent xy-locations, for each channel, then aligned and normalized (see Methods). The corresponding quantification of PRLR and E-cadherin fluorescence intensities across the plasma membrane indicates the relatively high intensity of intracellular PRLR signal compared with E-cadherin intensity peaking solely at the membrane (Figures 2B and 2D). While the intracellular localization pattern for PRLR is consistent with previous studies (Tallet et al., 2011; Lopez-Pulido et al., 2013; Millette et al., 2022), this pattern and its implications have not been further investigated. As pancreatic β-cells respond to glucose signaling, we perform glucose stimulation to consider any effect of additional physiological signals on PRLR localization behavior. Exposing live INS-1E cells and whole islets to low glucose (2.8 mM; to induce basal insulin secretion) and high glucose (16.7 mM; to induce stimulated insulin secretion) do not affect the tendency for PRLR to localize internally (Figures 2B and 2D). The similarity of the localization patterns, both qualitative and quantitative, across the two tissue types is striking and suggests that these patterns might have significance within PRLR signaling and organization. Consequently, intracellular localization should be observed in a reliable reconstruction model of PRLR expression.

**Figure 2:**
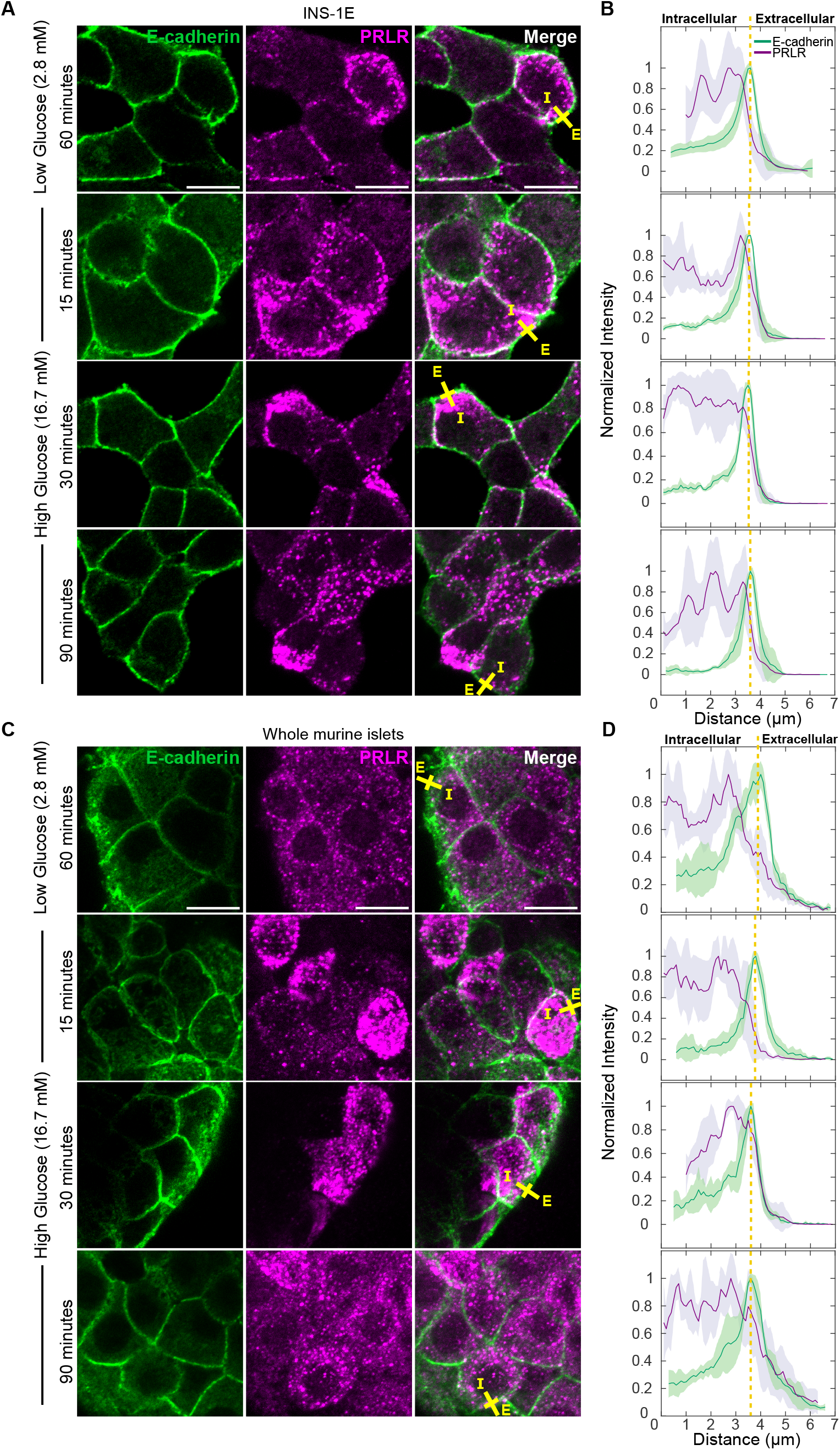
PRLR localizes heterogeneously under native expression in rodent tissue. **A** Representative confocal images of PRLR (shown in magenta) and E-cadherin (green) immunofluorescence in fixed INS-1E cells exposed to low glucose for 60 minutes and high glucose for 15, 30, and 90 minutes following 30 minutes of low glucose equilibration treatment. **B** Average fluorescence intensity is quantified along the representative yellow line (drawn from “I” – intracellular – to “E” – extracellular) spanning the plasma membrane. Fluorescence intensity line profiles are taken from at least 5 independent xy-locations in multiple cells, and shaded regions indicate a single standard deviation. **C** Representative confocal images of PRLR (magenta) and E-cadherin (green) immunofluorescence in fixed whole murine islets exposed to low glucose for 60 minutes and high glucose for 15, 30, and 90 minutes following the 30-minute low glucose treatment. **D** Average fluorescence intensity is quantified spanning the plasma membrane by measuring line profiles in 5 independent xy-locations in multiple cells. Scale bars 10 µm. Vertical dashed line on line profile plots indicates approximate location of the plasma membrane. Images are maximum intensity projections.

### Engineered minimal PRLR expression system reproduces endogenous PRLR localization pattern

To better understand PRLR localization patterns at the level of individual, live cells, and to build a system capable of studying both human and rodent PRLR, we develop a minimal expression system of human PRLR (hPRLR) in cells that do not otherwise express PRLR. Such an engineered system has been used to study pre-T cell receptor (preTCR) expression (James and Vale, 2012; Smid et al., 2023). We then apply our minimal engineered PRLR system to perform live fluorescence imaging. In this system, we label the extracellular domain of hPRLR with the constitutively fluorescent yellow-green fluorescent protein mEGFP. When expressed via transient transfection, this fluorescent protein fusion enables live imaging studies to investigate localization as well as molecular characteristics such as diffusion and oligomerization of hPRLR. As a control condition, we label CD86, a well-characterized monomeric type I transmembrane protein, with mEGFP. When transiently expressed in HEK293T cells, CD86-mEGFP is clearly observed at the plasma membrane (Figure 3A). Live confocal imaging of transient mEGFP-hPRLR expression in HEK cells in the minimal, transient expression system shows predominantly intracellular PRLR localization (Figure 3B), following a pattern similar to that observed in fixed rodent tissue (Figure 2). Quantifying the fluorescence intensity of each receptor as an xy-line profile indicates CD86 localization primarily at the plasma membrane and hPRLR localization distributed throughout the cytoplasm in addition to localization at the membrane (Figure 3C). We observe this localization pattern across cell lines (Supplementary Figure S1A) and when fusing different signal peptide sequences to hPRLR (Supplementary Figure S1B). To assess the effects of glucose stimulation on PRLR organization in the minimal system, we repeat live confocal imaging following treatment with glucose and do not observe a change in hPRLR localization (Supplementary Figure S1C). Thus, we go on to apply imaging and spectroscopy tools within our live engineered system to further investigate receptor organization and dynamics.

**Figure 3:**
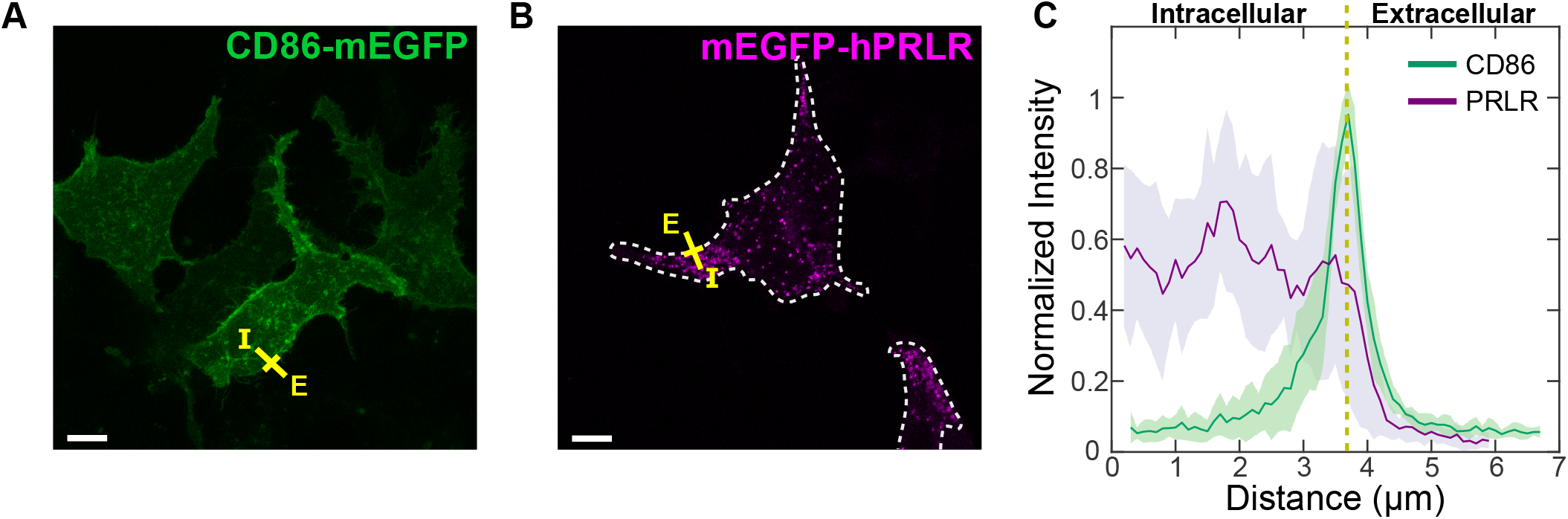
PRLR localizes heterogeneously in a minimal engineered expression system. **A** Representative confocal image of live membrane receptor expression for CD86-mEGFP (green) in HEK293T cells. **B** Representative confocal image of live membrane receptor expression for mEGFP-hPRLR (magenta) in HEK cells. Dotted line indicates location of cell membrane. **C** Average fluorescence intensity is quantified along the representative yellow line (drawn from “I” – intracellular – to “E” – extracellular) spanning the plasma membrane. Fluorescence intensity line profiles are taken from 10 independent xy-locations in multiple cells, and shaded regions indicate a single standard deviation. Vertical dashed line on line profile plot indicates approximate location of the plasma membrane. Scale bars 10 µm. Images are maximum intensity projections.

### Live imaging and spectroscopy characterize PRLR populations in minimal expression system

We next set out to visually and quantitatively distinguish cell surface-localized and intracellular-localized populations of hPRLR in our reconstituted system. First, to determine whether we can visualize these two populations of hPRLR, we fuse a FLAG sequence (DYKDDDDK) to the extracellular domain of our mEGFP-hPRLR construct. We target the FLAG sequence in a second fluorescent channel using commercially available anti-FLAG antibodies in non-permeabilized live and fixed cells. By adding fluorescently labeled anti-FLAG antibodies to non-permeabilized cells, followed by confocal imaging, we can specifically target membrane-localized hPRLR (Figure 4A). Live imaging of anti-FLAG labeling suggests rapid internalization of hPRLR, while live and fixed anti-FLAG visualizations show two distinct PRLR populations – one at the plasma membrane, and one that is intracellular (Figure 4B).

**Figure 4:**
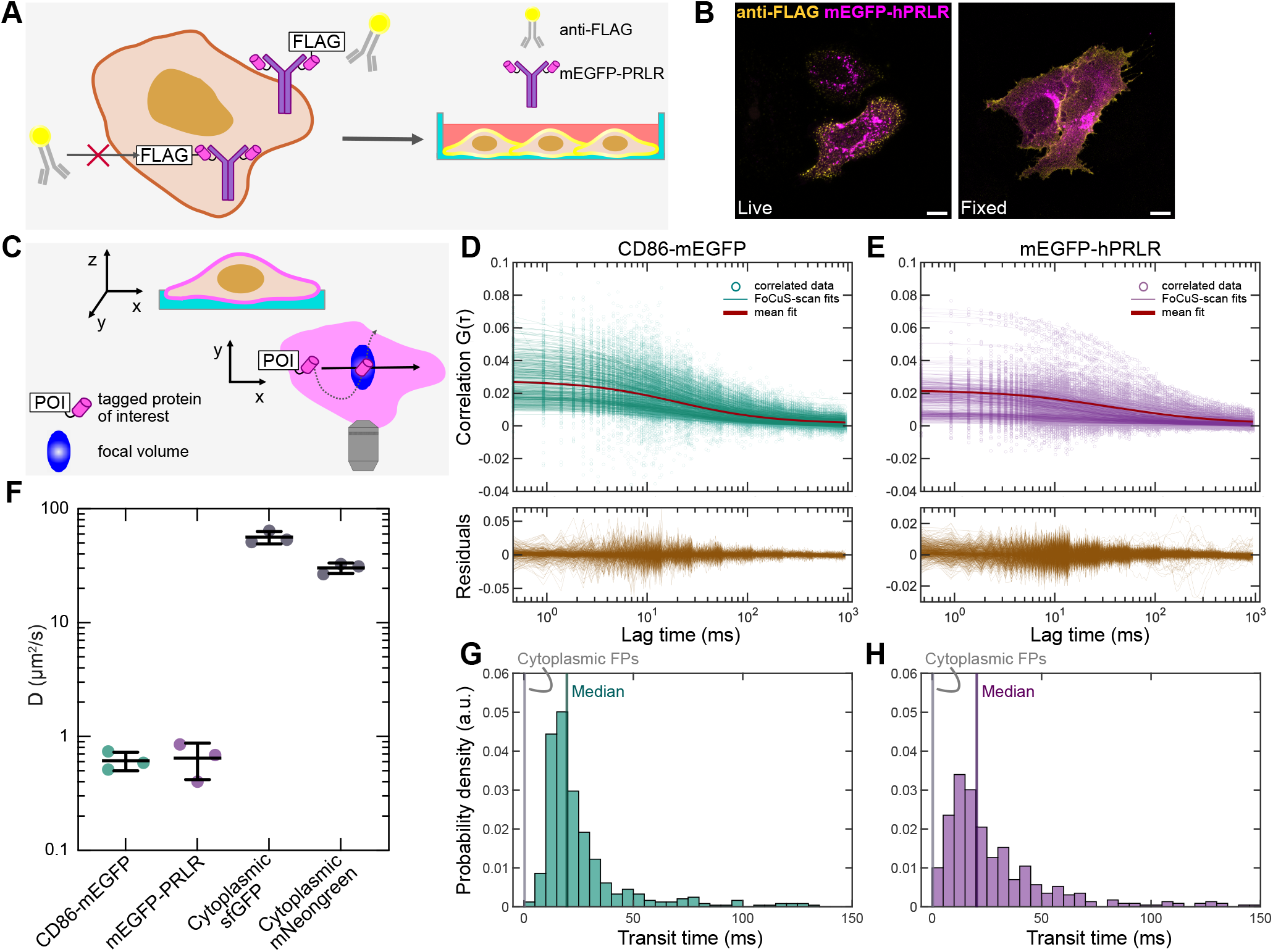
Live imaging and spectroscopy spatially resolve cell surface- and intracellular-localized hPRLR populations. **A** Schematic of anti-FLAG labeling to target membrane-localized PRLR in non-permeabilized cells. **B** mEGFP-hPRLR (magenta) and FLAG expression (yellow) in live and fixed U2OS cells. Scale bars 10 µm. **C** Schematic of scanning FCS (sFCS) acquisition in the basal membrane of live cells. FCS acquisitions measure fluctuations in fluorescence intensity as a fluorescent-labeled protein of interest diffuses in and out of the focal volume of a confocal microscope. sFCS measurements are acquired at multiple points along a scanned line within the sample. **D, E** sFCS fitting using residuals between correlated data and FoCuS-scan models fit to photophysical parameters for CD86-mEGFP and mEGFP-hPRLR. **F** Diffusion coefficients measured for three replicates each of CD86-mEGFP, mEGFP-hPRLR, and cytoplasmic mNeonGreen and sfGFP. Cytoplasmic fluorescent proteins are measured using point-FCS. **G, H** Transit times calculated from fitting for CD86-mEGFP and mEGFP-hPRLR. Median transit times are displayed as a vertical line for each protein.

To quantitatively probe the dynamics of membrane-localized hPRLR, we use scanning fluorescence correlation spectroscopy (sFCS) to measure diffusion coefficients, *D*, of mEGFP-hPRLR in the basal membrane of transfected cells (Figure 4C). Quantifying diffusion provides context for the behavior and ultimately the spatial context of the measured protein. CD86-mEGFP is again used as a control for sFCS measurements. As an additional control condition, we measure cytoplasmic diffusion of mNeonGreen and sfGFP using point-FCS. Because sFCS measures molecular diffusion along a scanned line, this method best captures slow and spatially heterogeneous dynamics typical of protein movement within a lipid membrane (Levi et al., 2003). In contrast, point-FCS is performed at a single diffraction-limited spot and best captures fast dynamics in a more homogeneous molecular environment. Correlation curves are fit to photophysical models using FoCuS-scan for sFCS measurements (Waithe et al., 2018) and FoCuS-fit-JS (Waithe, 2021) for point-FCS measurements. Correlation curve fits indicate the presence of a single species diffusing slowly for the CD86 control (Figure 4D) and hPRLR (Figure 4E). We measure D = 0.61 ± 0.11 µm^2^/s for CD86 and D = 0.65 ± 0.23 µm^2^ /s for hPRLR (Figure 4F). Our measured *D* value for CD86 is consistent with reported literature values for type I transmembrane receptors (Gakamsky et al., 2005; Baker et al., 2007; Schneider et al., 2024). We measure D = 55.9 ± 6.6 µm^2^ /s for cytoplasmic sfGFP and D = 30.5 ± 4.1 µm^2^ /s for cytoplasmic mNeonGreen (Figure 4F). From fitting, the corresponding median transit times for the membrane proteins are calculated to be τ_D_ = 27.3 ms for CD86 (Figure 4G) and τ_D_ = 23.5 ms for hPRLR (Figure 4H). Additional sFCS replicates are shown in Supplementary Figure S2. We observed median transit times of τ_D_ 0.26 ms and τ_D_ = 0.52 ms for cytoplasmic sfGFP and mNeonGreen, respectively. Previously reported values for diffusion of a single-pass transmembrane protein are approximately 0.5 µm^2^ /s (Schneider et al., 2024), while cytoplasmic diffusion is measured on the order of 10^0^ to 10^1^ µm^2^ /s (Mørch and Schneider, 2023). Together with the measured diffusion coefficients, the measured transit time values for both membrane receptors are consistent with membrane diffusion. Measuring PRLR diffusion on the scale of CD86 diffusion confirms that PRLR is present at the plasma membrane.

### Simulations of trafficking model identify parameter sensitivities

To investigate the implications of intracellular PRLR localization across timescales in the context of JAK/STAT signaling in β-cells, we include PRLR trafficking within an existing mathematical model of the signaling pathway (Mortlock et al., 2021) (Figure 5A). This cycling mechanism is known to occur in other signaling pathways. For example, vascular endothelial growth factor (VEGF) receptors have been shown to traffic between the cell surface and intracellular compartments (Simons, 2012), and existing computational models of VEGF signaling describe these receptor trafficking dynamics (Tan et al., 2013; Clegg and Gabhann, 2015). We expand the existing model of PRLR-mediated signaling such that in addition to participating in signal transduction reactions at the cell surface, PRLR complexes are internalized within an intracellular compartment and recycled from that compartment back to the cell surface. Within the intracellular compartment, internalized PRLR complexes (denoted with “i” in the schematic) participate in the same set of reactions that occur at the cell surface, with the exception of the forward PRL binding reaction. Our revised model uses the set of parameters and initial species concentrations with the lowest error between model predictions and experimental data found by Simoni et al. (Simoni et al., 2022).

**Figure 5:**
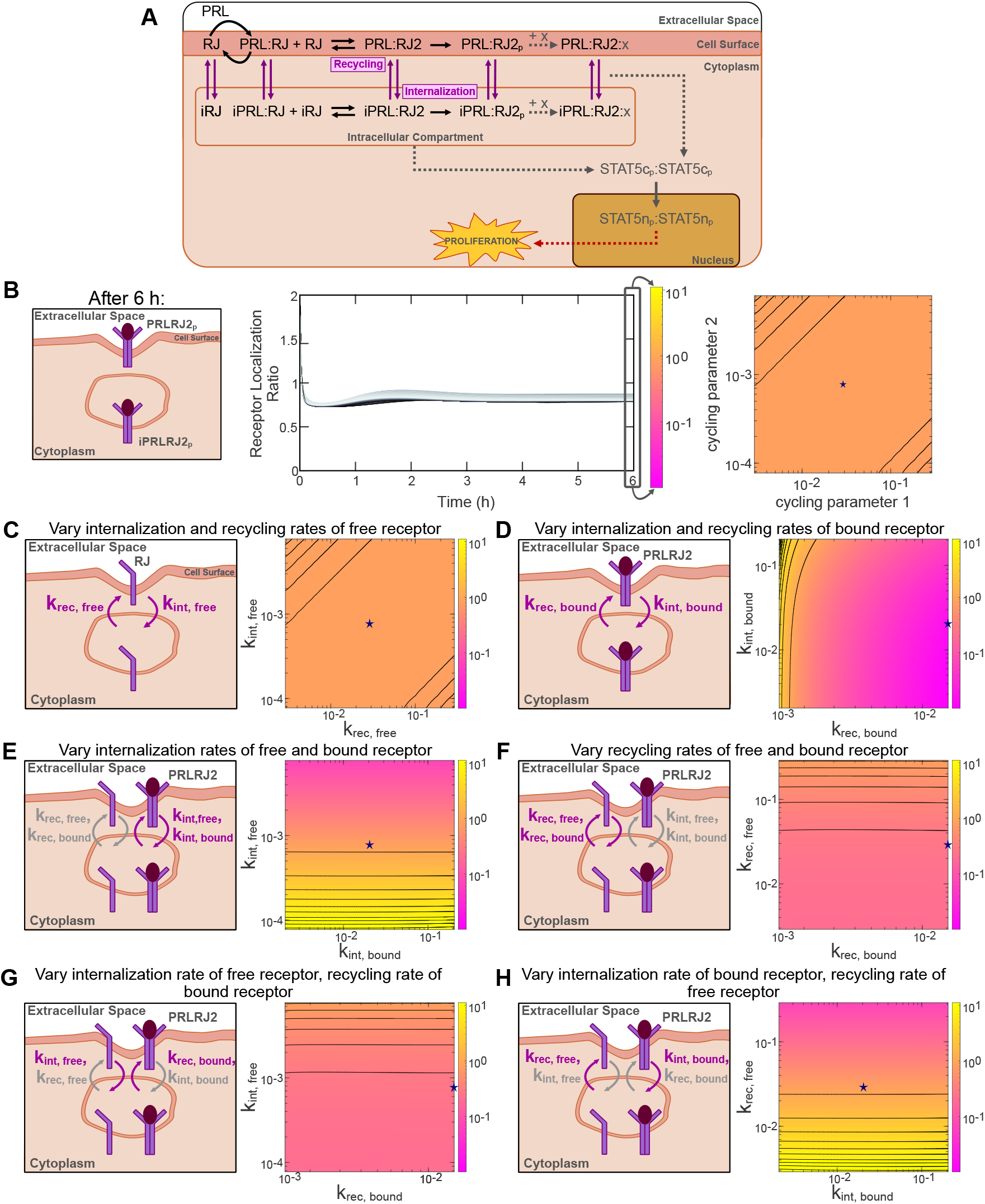
Integrating intracellular PRLR trafficking and exploring the trafficking parameter space. **A** Schematic of ODE model reactions in the updated trafficking model. The reactions in the intracellular compartment, and the internalization of cell-surface PRLR complexes and recycling of intracellular PRLR complexes to the cell surface, build on the model created by Mortlock et al. (Mortlock et al., 2021). Species that participate in reactions in the intracellular compartment are denoted with “i” preceding the species name. “x” indicates reactants that can form complexes with PRLR. RJ and iRJ are the receptor starting material in the respective compartments. **B** Each pair of internalization/recycling rates is used in independent simulations of the trafficking model to investigate activated receptor (PRLRJ2_p_, iPRLRJ2_p_) localization. Each rate is sampled within an order of magnitude below and above its baseline value. The ratio of active receptor dimers at the cell surface to active receptor dimers within the intracellular compartment following 6 h of simulated time is visualized as a 2D surface. Contours are shown as black curves. The coordinate pair consisting of the two baseline values is denoted by a star. **C-H** Schematic representations of parameter combinations and corresponding 2D surfaces for varying recycling and internalization rates of free receptor, recycling and internalization rates of bound receptor, the internalization rate of free and bound receptor, the recycling rate of free and bound receptor, internalization rate of free receptor and recycling rate of bound receptor, and internalization rate of bound receptor and recycling rate of free receptor. All results shown are for the initial condition of all RJ localized internally. Purple arrows indicate the rate or rates being varied in each condition. Gray arrows indicate the rate or rates being held constant at baseline. k_rec, bound_ is only varied below the value reported by Tan et al. due to ODE solver tolerances.

We first asked how sensitive the revised PRLR trafficking model is to varying the trafficking rates of PRLR complexes. To explore the possible parameter space describing PRLR trafficking, we consider and vary four trafficking rates governing movement of PRLR between the cell surface and intracellular compartments: (1) k_int, bound_, the internalization rate of ligand-bound PRLR complexes, (2) k_rec, bound_, the recycling rate of ligand-bound PRLR complexes, (3) k_int, free_, the internalization rate of non-ligand bound (free) PRLR complexes, and (4) k_rec, free_, the recycling rate of free PRLR complexes. We create a coordinate grid covering the search space and simulate the trafficking model using each coordinate pair. All baseline trafficking rate values are taken from previous work on VEGF trafficking by Tan et al. (Tan et al., 2013). We vary the previously reported rates within an order of magnitude below and above the baseline value, with the exception of the recycling rate of ligand-bound PRLR (k_rec, bound_), which is only varied below the baseline value due to ODE solver tolerances.

To assess the effects of PRLR trafficking on JAK/STAT signal transduction, we ask whether the majority of activated receptor complexes are present at the cell surface or localized internally 6 h post-PRL stimulation (Figure 5B). This result is visualized as a heatmap relating the ratio of surface-(PRLRJ2_p_) to intracellular-localized (iPRLRJ2_p_) activated receptor, or receptor localization ratio. We vary the previously fit value of the receptor starting material, RJ (Figure 5A), under five independent initial conditions: RJ is initialized with 100%, 75%, 50%, 25%, or 0% of the fit value within the intracellular compartment, and the remaining amount is initialized at the cell surface. We specifically vary the localization of PRLR starting material, rather than investigate values for the absolute amount of RJ and iRJ, to apply and better understand our experimental findings. All results shown are simulated with the initial condition of 100% intracellular-localized RJ, as the initial distribution does not qualitatively affect the model simulation outputs (Supplementary Figure S3A), and our imaging data suggest a relatively large intracellular receptor pool. We observe that varying the trafficking rates for free receptor always leads to a greater amount of intracellular-localized activated receptor than surface-localized, suggesting the intracellular forms of PRLR act as a sink (Figure 5C). For all other trafficking rate pairs, there are combinations of values that lead to a ratio of surface-to-internal localized activated receptor greater than 1 (Figure 5D-H). The level of surface-localized activated receptor is maximized by increasing the internalization rate of bound receptor at low rates of free receptor trafficking (Figures 5E and 5H).

To investigate the implications of intracellular PRLR’s ability to promote signaling in our model, we then individually varied the rate of three key reactions in promoting and inhibiting signal transduction. We vary the rate at which activated, intracellular PRLR-JAK dimers phosphorylate cytosolic STAT5 monomers (model parameter k6) to study the ability of intracellular PRLR to activate STAT5. We vary the rate at which activated, intracellular PRLR-JAK dimers are inactivated by SHP-2 (k10) to simulate how dephosphorylation of intracellular PRLR bound to JAK influences signaling. Finally, we vary the rate at which activated, intracellular PRLR-JAK dimers are inactivated by suppressor of cytokine signaling (SOCS1; k21) to investigate the impact of negative feedback. These rates are varied only in equations representing reactions that occur within the intracellular compartment, namely, each rate (k6, k10, and k21) takes on the best fitted value from our prior work (Simoni et al., 2022; Huber et al., 2023) for all reactions occurring at the cell surface, and three additional parameters are introduced and used for reactions occurring within the intracellular compartment (k6_internal_, k10_internal_, and k21_internal_). We model the effects of varying these rates on four downstream signaling outputs: (1) the ratio of nuclear/cytosolic STAT5A up to 6 h post-PRL stimulation, (2) the ratio of nuclear/cytosolic STAT5B up to 6 h post-PRL stimulation, (3) phosphorylated JAK (pJAK) dimers up to 6 h post-PRL stimulation, and (4) the fold change of Bcl-xL up to 24 h post-PRL stimulation. These four outputs represent critical signaling species that influence β-cell function. We again vary these three kinetic rates under five independent sets of initial conditions describing the distribution of PRLR starting material, species RJ: 100%, 75%, 50%, 25%, or 0% intracellular. All results shown are for the initial condition of all RJ localized internally. Interestingly, we do not observe a difference in sensitivity of any outputs when we vary the initial localization of RJ (Supplementary Figure S3B). This suggests that our model of this pathway is more robust to changes in initial conditions than to changes in individual reaction rates and that the model equilibrates quickly. Additionally, we find that downstream readouts are more sensitive to varying how intracellular PRLR mediates phosphorylation of STAT5 (k6_internal_, Figure 6A) than to varying the inhibitory reactions (k10_internal_ and k21_internal_, Figures 6B and 6C). Thus, the ability of intracellular PRLR to mediate STAT5 activation strongly influences downstream signaling.

**Figure 6:**
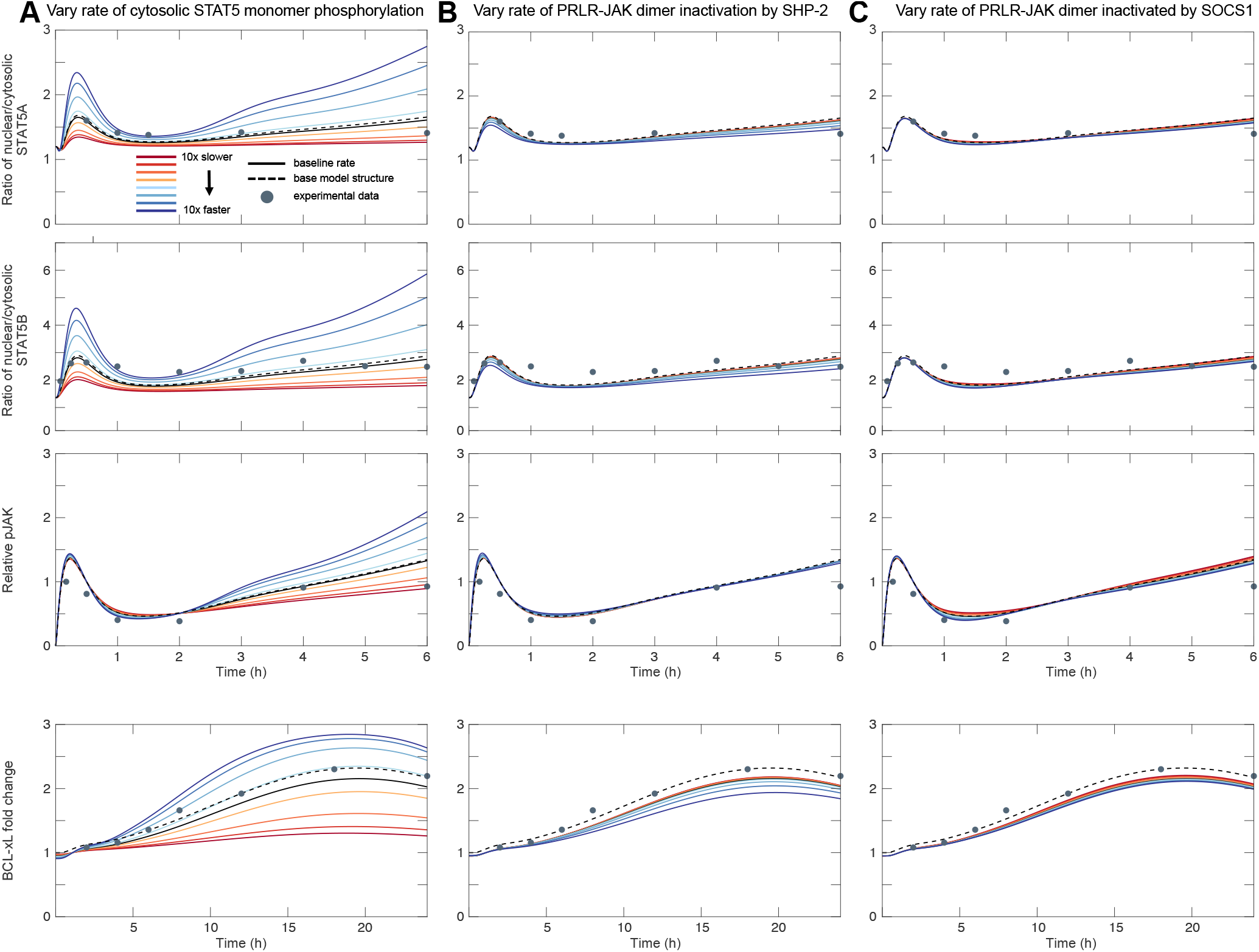
PRLR signaling readouts are sensitive to varying forward signal transduction. **A-C** Responses to varying the rate of cytosolic STAT5 monomer phosphorylation by active, intracellular PRLR-JAK dimers (k6_internal_), intracellular PRLR-JAK dimer inactivation by SHP-2 (k10_internal_), and intracellular PRLR-JAK dimer inactivation by SOCS1 (k21_internal_) within one order of magnitude below and above the best fitted parameter value for each rate. All results shown are for the initial condition of all RJ localized internally. Simulations using the baseline rates are shown by solid black curves. Simulations using the base model structure (without trafficking reactions) are shown by the dashed black curves, and experimental data (from Brelje et al. (Brelje et al., 2002)) are shown as points. STAT5 and pJAK activity is simulated over 6 hours, and BCL-xL expression is simulated over 24 hours.

## Discussion

Developing mathematical models fit to biological data demand active interaction between quantitative experimental data and the models themselves. Collectively, an integrated experimental-computational approach shapes our understanding of complex, multiscale systems such as signal transduction. Here, we design and implement the first steps of such an approach to study PRLR-mediated JAK/STAT signaling in pancreatic β-cells. We apply live imaging and powerful spectroscopic tools in combination with mathematical modeling to better understand the behavior of PRLR on the scale of individual cells. We build on previous studies that have reported intracellular PRLR localization by developing an integrated approach capable of asking what implications this localization pattern has on cell signaling. The utility of this approach is therefore two-fold, moving towards biological insights into a multiscale biological problem – here, PRLR localization in the context of β-cell signaling – and leveraging imaging tools with modeling to arrive at those insights.

Overall, we demonstrate that (1) PRLR exhibits heterogeneous spatial organization in primary pancreatic tissue (Figure 2), (2) a reconstituted minimal expression system of PRLR-fluorescent protein fusions reproduces PRLR localization patterns observed in primary tissue (Figure 3), (3) two spatially distinct PRLR populations are observed simultaneously in live and fixed cells, and those spatially distinct PRLR populations can be measured and characterized using FCS (Figure 4), (4) our experimental observations can be leveraged to incorporate receptor trafficking into a model of PRLR-mediated JAK/STAT signaling (Figures 5 and 6), and (5) PRLR-mediated JAK/STAT signaling is predicted to be more sensitive to varying reaction rates than to changes in the initial distribution of PRLR (Figures 5 and 6).

Our findings have biological implications for PRLR signaling. First, the finding that PRLR localizes internally in both its endogenous expression system and in an engineered system suggests the significance of a storage mechanism. Employing this storage mechanism could suggest that β-cells exhibit robustness to stimulation by external signals, which could play a role in modulating expression of pro-proliferative factors, thereby regulating populations of insulin-secreting cells. Our model predictions indicate that strategies to enhance reaction kinetics would have a greater impact on PRLR-mediated signaling compared with altering the localization pattern of surface and intracellular PRLR. Receptor trafficking leading to cell signaling rather than attenuation has been well studied in receptor tyrosine kinases (RTKs) (Wang et al., 2002; Horowitz and Seerapu, 2012) and there is continued interest in the field to elucidate the (patho)physiology of receptor trafficking fates across cell types (Cullen and Steinberg, 2018). More recently extensive work has been done to better understand the role of receptor trafficking in T-cell development using a reconstitution approach for pre-T cell receptor (preTCR) expression similar to the work presented here (James and Vale, 2012; Smid et al., 2023). Intracellular signaling networks in the context of pancreatic β-cell proliferation have been well mapped in rodent studies, though an understanding of the complex crosstalk between signals leading to human β-cell survival and proliferation is limited (Stewart et al., 2015; Jiang et al., 2018), thus motivating the development of tools tailored to answer questions regarding human β-cell signaling. Together, approaches based in a modeling-imaging framework developed for human β-cell studies can be applied to develop a mechanistic understanding of signaling dynamics in other physiological contexts.

We acknowledge the potential limitations of our study. We use fixed tissue immunofluorescence, which can in some cases alter protein expression patterns, though typically this affects DNA-protein interactions (Irgen-Gioro et al., 2022; Yoshida et al., 2023). Transgenic mice expressing fluorescently labeled PRLR signaling proteins could be used to expand the study of endogenous PRLR dynamics. Here we only show the behavior of labeled PRLR. An expansion of the reconstituted system could label STAT5 in addition to PRLR to track the intracellular effects of PRLR stimulation at the plasma membrane across spatiotemporal scales. Multiscale analyses could also be extended to study the nanoscale changes in PRLR organization. Here we apply FCS to spatially resolve PRLR localization; FCS is also suited for oligomerization studies. Applied to this system, FCS could be used to ask questions about what changes in PRLR’s behavior occur upon dimerization or oligomerization. To do this, we can use both anti-PRLR and anti-FLAG antibodies as well as physiological signals, such as prolactin, to induce oligomerization (Digman et al., 2008; Schneider et al., 2024). Further, constitutive fluorescence poses a limitation to resolving the temporal dynamics of PRLR trafficking. An interesting approach could replace the fluorescent protein tag with a chemigenetic label such as SNAP-tag to quantify PRLR internalization from the cell surface by using a membrane-impermeable substrate. Such trafficking data can be used to further refine the parameter space for models of PRLR signaling. Addressing these questions will contribute to efforts to understand how PRLR signaling is integrated to lead to pro-proliferative gene expression in pancreatic β-cells. Our present experimental approach lays a framework for such work towards elucidating potential therapeutic mechanisms within signal transduction networks. Finally, the revised mathematical model can be further explored by performing an extended Fourier Amplitude Sensitivity Test (eFAST) (Marino et al., 2008) and by investigating the assumption that internalized PRLR species can participate fully in signaling. Global sensitivity analysis of model parameters performed by eFAST would determine if the kinetic rates describing trafficking reactions significantly impact the model predictions.

Altogether, we use a minimal system reconstitution approach to further investigate prior findings of intracellular PRLR localization, and we translate our experimental findings into a mathematical representation of receptor trafficking within a revised model of PRLR-mediated JAK/STAT signaling. We find that a reconstitution approach readily reproduces endogenous PRLR localization patterns in a minimal system, thus producing a valuable new tool for further study of PRLR dynamics in the context of β-cell signaling. We identify two spatially distinct PRLR populations in our reconstituted system. Finally, when we leverage evidence of those populations in a mathematical model, model predictions indicate higher sensitivity to trafficking reactions rates than to the initial spatial distribution of PRLR. In conclusion, quantitative imaging in combination with mathematical modeling provides a powerful, complementary tool for revealing molecular dynamics in cell signaling networks.

## Methods

### Plasmid design

A plasmid containing human prolactin receptor (hPRLR), fused to the rat PRLR signal peptide, was obtained as a kind gift from Vincent Goffin (INSERM) (Bogorad et al., 2008). The pcDNA3 FLAG HA plasmid was obtained as a gift from William Sellers (Addgene plasmid #10792; http://n2t.net/addgene:10792; RRID:Addgene_10792). A signal peptide (rat PRLR, human PRLR, or human CD86 signal peptide), hPRLR, and a yellow-green fluorescent protein (mNeonGreen, and, separately mEGFP) are cloned into the pcDNA3 plasmid using Takara In-Fusion Snap Assembly (#638947) according to the manufacturer’s protocol. We create two plasmid designs, fusing each fluorescent protein to (1) the N-terminus of hPRLR, to create a fusion of the fluorescent protein at the extracellular domain of hPRLR, and (2) the C-terminus of hPRLR, to create a separate, intracellular fusion. CD86 is fused with mEGFP as described above. The CD86 sequence was obtained as a gift from Alexander Mørch (University of Oxford).

### Tissue culture

U2OS and HEK293T cells were acquired from the American Type Culture Collection. U2OS and HEK cells are cultured in DMEM (Corning, #10-013-CV), supplemented with 10% fetal bovine serum (FBS; Thermo Fisher Scientific, #A5256701), 1% penicillin/streptomycin (Thermo Fisher Scientific, #15140122), and 1% L-glutamine (Sigma-Aldrich, #G7513). INS-1E cells were acquired from Pierre Maechler (Université de Genève) and are cultured in optimized RPMI-1640 medium (AddexBio, #C0004-02). Murine islets were provided by Kate White (USC).

Prior to imaging experiments, cells are grown to approximately 80% confluency before being seeded onto 8-well #1.5 glass chambered coverslips (ibidi, #80807). 24 h after seeding, transient transfection of pcDNA3-hPRLR plasmids is performed in HEK or U2OS cells using Lipofectamine 3000 and P3000 reagents (Thermo Fisher Scientific, #L30008) according to the manufacturer’s protocol. 24 h after transfection, cell culture medium is replaced with phenol red-free Leibovitz’s L-15 medium (Thermo Fisher Scientific, #21083027) for imaging.

### Glucose treatment and immunofluorescence

To validate PRLR localization patterns in native tissue, immunofluorescence (IF) is performed on INS-1E cells and whole murine islets following exposure to low and high glucose solutions. For experiments involving primary β-cells, animal procedures were approved and conducted per the IACUC guidelines at the University of Southern California under Animal Use Protocol #21120. Glucose treatment is performed, without fixation, on transiently transfected U2OS cells. IF is also performed in fixed and live transiently transfected U2OS cells to visualize hPRLR populations using anti-FLAG antibodies.

To begin insulin secretion experiments, INS-1E cells, whole islets, and transiently transfected U2OS cells are starved in low (2.8 mM) glucose solution (HEPES-buffered Krebs-Ringer buffer, pH 7.4, 0.1% bovine serum albumin (BSA), 111 mM NaCl, 25 mM NaHCO_3_, 4.8 mM KCl, 1.2 mM KH_2_PO_4_, 1.2 mM MgSO_4_, 10 mM HEPES, 2.3 mM CaCl_2_) for 30 minutes at 37°C. For basal insulin secretion, cells and islets are then incubated in fresh 2.8 mM glucose solution for 1 h at 37°C. For stimulated insulin secretion, cells and islets are instead incubated in high (16.7 mM) glucose solution for 15, 30, 45, 60, and 90 min. Transfected U2OS cells are imaged live following the respective glucose treatments.

INS-1E cells are fixed in 4% paraformaldehyde (PFA) for 15 min at room temperature, and whole islets are fixed in 4% PFA overnight on a shaker at 4°C. 8-well chambered coverslips are prepared for adhering fixed islets by coating with poly-D-lysine (PDL; Thermo Fisher Scientific, #A3890401) for 24 h at 4°C. Islets are then plated and allowed to adhere overnight at 4°C in 1X phosphate buffered saline (PBS), on PDL-coated slides. For glucose experiments, INS-1E cells and islets are permeabilized prior to antibody incubation. Permeabilization, where indicated, is performed using 0.1% Triton X-100 in PBS at room temperature for 10 minutes. Prior to incubating primary antibodies, a blocking buffer of 5% FBS, 2% BSA in PBS is applied for 1 h at room temperature. INS-1E cells and islets are stained with anti-PRLR (Bioss, #BS-6445R) and anti-E-cadherin (R&D Systems, #AF748). Fixed and live U2OS cells are stained with anti-FLAG (BioLegend, #637319). Anti-PRLR and anti-E-cadherin are diluted 1:200 in 1% BSA and incubated overnight at 4°C. Fluorescent secondary antibodies are diluted 1:1000 in PBS and incubated for 30 min at room temperature prior to imaging.

For anti-FLAG experiments, live and fixed U2OS cells are not permeabilized prior to antibody incubation. In fixed cells: 24 h after transfection, cells are fixed in 4% PFA for 15 min at room temperature and blocked with 5% FBS, 2% BSA in PBS for 1 h. Anti-FLAG is diluted 1:200 in 1% BSA and incubated overnight at 4°C. In live cells: 24 h after transfection, anti-FLAG is diluted 1:200 in DMEM and incubated for 30 min at 37°C. Fluorescent secondary antibodies are diluted 1:1000 in PBS and incubated for 30 min prior to imaging.

### Confocal fluorescence microscopy

All fluorescence imaging is performed on a Zeiss LSM 880 confocal microscope with a Zeiss LD C-Apochromat 40x 1.1 NA water objective. mNeonGreen, mEGFP, and sfGFP are excited using the 488 nm laser line and 488/561/633 beam splitter, and emission is detected between 500 and 600 nm with the ChS1 detector. Live imaging is performed at 37°C.

### Fluorescence intensity line profile analysis

Fluorescence intensity line profiles are measured in FIJI (Schindelin et al., 2012). Lines are each 1 pixel in width and approximately 5 µm in length. Each line is centered on the plasma membrane using either the fluorescent channel or transmitted light (T-PMT). Lines are sampled at independent locations in multiple cells for each condition. MATLAB’s (MathWorks, R2024a) xcorr function is used to align repetitions of each visualized protein and condition. Intensities are normalized to the maximum intensity value for each condition.

### FCS data acquisition and analysis

Point and scanning fluorescence correlation spectroscopy (point- and sFCS) measurements are acquired on a Zeiss LSM 880 confocal microscope using the FCS mode (point measurements) and xt line scanning (scanning measurements) included in Zeiss’ Zen Black software.

All measurements are taken in live cells at 37°C as described previously (Schneider et al., 2021; Mørch and Schneider, 2023). Briefly, measurements are first calibrated using a solution of 50 nM Alexa Fluor 488 in water by adjusting the objective’s correction collar and aligning the pinhole. Following calibration, 10-second repetitions of point-FCS measurements are acquired at individual locations within the cytoplasm of live cells and saved as.fcs files. sFCS measurements are acquired at individual locations within the plasma membrane (for CD86 and hPRLR experiments) as 52-pixel xt line scans at approximately 2000 Hz with a pixel dwell time of 3.94 µs. Line scans are saved as.lsm files.

FCS data are analyzed using the Python-based, open-source FoCuS-scan software (Waithe et al., 2018). Here, FoCuS-scan is run in a semi-custom Python environment on macOS Monterey 12.5.1, with the packages detailed at https://github.com/lynne-cherchia/sFCS-macOS. For curve fitting, we assume a value of 1 for sub-diffusion parameter *α* and a triplet time tauT1 of 0.005 ms for Alexa Fluor 488 calibration measurements and 0.04 ms for fluorescent protein measurements. For diffusion dimensionality, we fit free-dye calibration and cytoplasmic fluorescent protein measurements in 3D and restrict diffusion to 2D for hPRLR and CD86 measurements. For photobleaching correction in sFCS measurements, we crop the first 5-10 seconds and apply a local averaging correction (Waithe et al., 2018).

### Base JAK/STAT model and updated trafficking model structure

An ordinary differential equation (ODE) model describing the kinetics of JAK/STAT signaling activated by PRLR in pancreatic β-cells was previously constructed in MATLAB (Mortlock et al., 2021). The model uses ODEs to describe the reactions involved in intracellular PRLR signaling: Prolactin (PRL) binds to PRLR, inducing receptor dimerization and subsequent activation of JAK2 and STAT5 by phosphorylation. Activated STAT5 monomers then dimerize in the cytosol, translocate to the nucleus, and finally act as transcription factors promoting transcription and translation of target genes, including PRLR, suppressors of cytokine signaling (SOCS), and the anti-apoptotic protein B-cell lymphoma-extra large (Bcl-xL). The model comprises 56 species, and 56 corresponding ODEs describing their time-dependent dynamics, and 61 parameters. Bayesian inference was used previously to estimate the kinetic parameters from experimental data found in the literature (Simoni et al., 2022); because the focus of applying the model in this study is to investigate a specific biological question, we only use the parameter set with the lowest error between model predictions and experimental data.

To reflect our experimental findings in the existing model, we add 11 equations and 4 parameters describing internalization and recycling of activated PRLR, effectively adding a compartment to the model structure by distinguishing between membrane-localized and intracellular PRLR (Figure 5A). We add 11 species, with 11 corresponding ODEs, to account for all forms of the receptor complex expressed either in a “pool” at the plasma membrane or in an intracellular “pool.” It is also likely that PRLR is cycled between these two pools. To describe this movement, we add 4 parameters: (1) k_int, bound_, the internalization rate of ligand-bound PRLR complexes, (2) k_rec, bound_, the recycling rate of ligand-bound PRLR complexes, (3) k_int, free_, the internalization rate of non-ligand bound (free) PRLR complexes, and (4) k_rec, free_, the recycling rate of free PRLR complexes. This cycling mechanism is known to occur in VEGF signaling (Simons, 2012) and has previously been implemented in a similarly structured ODE model of VEGF signaling (Tan et al., 2013). The reaction network is detailed in Supplementary Table S1. Here, we compare simulation results predicted by (1) the original model developed by Mortlock et al. (Mortlock et al., 2021) and (2) the updated version of the model that reflects our experimental findings. All of the code necessary to simulate the trafficking model is available at https://github.com/lynne-cherchia/PRLR-trafficking.

### Simulating variable initial PRLR distribution

We simulate the effects of our imaging results by varying the initial location of PRLR. For all model simulations, we consider the following initial distributions of plasma membrane- and intracellular-localized PRLR starting material, species RJ (and iRJ), in the updated trafficking model: 100%, 75%, 50%, 25%, and 0% intracellular.

### Cycling parameter space exploration

To begin exploring PRLR’s potential cycling dynamics, we use rate values from an established model of VEGF signaling (Tan et al., 2013). We explore the pairwise effects of varying these four rates – internalization and recycling of both free (k_int, free_, k_rec, free_) and ligand-bound receptor (k_int, bound_, k_rec, bound_) – on the localization of active receptor dimers. We vary the four baseline cycling parameters within an order of magnitude below and above the values given by Tan et al. and use MATLAB’s meshgrid function to create a 250-by-250 grid corresponding with parameter value coordinates for each pairwise combination of the four cycling rates. The recycling rate of ligand-bound PRLR (k_rec, bound_) is only varied below the reported baseline value due to failure of the solver to converge for rates above baseline. ODEs are solved using the MATLAB ode15s solver. Using high-performance computing provided by the USC Center for Advanced Research Computing, we then simulate the compartment model with these varied cycling rates and explore the effects on the receptor localization ratio – the ratio of cell surface-localized activated PRLR dimers (PRLRJ2_p_) to intracellular-localized activated PRLR dimers (iPRLRJ2_p_) after 6 h of simulated time.

### Sensitivity analysis of intracellular PRLR

We perform a sensitivity analysis to evaluate how model predictions of key outputs are affected by the presence of intracellular PRLR. To test the effects of intracellular PRLR complexes exhibiting varied ability to transduce signal, we vary the following kinetic rates, within the intracellular compartment only, from their baseline values: (1) k6_internal_, the rate at which activated, intracellular PRLR-JAK dimers phosphorylate cytosolic STAT5 monomers, (2) k10_internal_, the rate at which activated, intracellular PRLR-JAK dimers are inactivated by SHP-2, and (3) k21_internal_, the rate at which activated, intracellular PRLR-JAK dimers are inactivated by SOCS1. Each rate is varied within an order of magnitude below and above the lowest error rates found by Simoni et al. (Simoni et al., 2022) for k6, k10, and k21. For each rate pair that is varied, all other kinetic rates are held constant. We model the effects of varying PRLR’s ability to transduce signal on STAT5 localization, pJAK levels, and Bcl-xL production.

## Acknowledgements

We thank Kate White, Senta Georgia, Janielle Cuala, and Zhongying Wang for providing mouse islets and for helpful discussions on rodent tissue experiments, Arkadi Shwartz and Jason A. Junge for managing the Translational Imaging Center, and the USC Tissue Culture Core for providing INS-1E cells. We also thank Dr. Vincent Goffin for the hPRLR plasmid. We thank members of the Finley research group for constructive feedback on the manuscript. L.C. acknowledges funding from a USC Graduate School Provost Fellowship. F.S. acknowledges funding from EMBO (EMBO ALTF 849-2020), HFSP (LT000404/2021-L), and Leverhulme Turst (LIP-2021-017).

