## Supplementary Material for "Prolactin receptor localization and dynamics: Insights from quantitative imaging and mathematical modeling"

### Supplemental Material

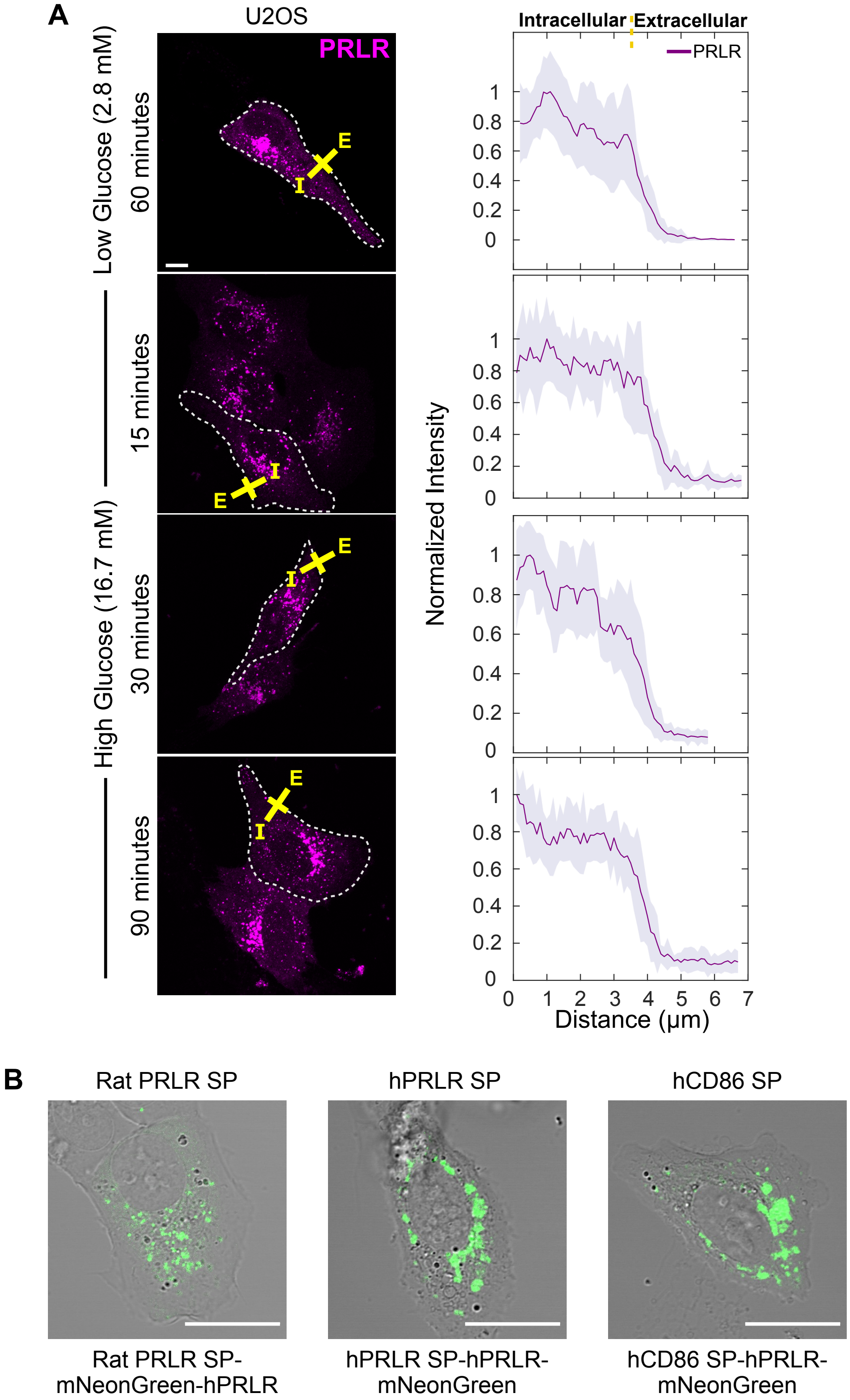


**Figure S1: Intracellular hPRLR localization is observed across cell lines and when the signal peptide is varied. A** Intracellular hPRLR is seen when the mEGFP-hPRLR fusion is expressed in a different cell line (here, U2OS). Glucose stimulation experiment is repeated in the minimal system and as expected, hPRLR localization is unchanged when exposed to low and high glucose solutions. Dotted line indicates location of cell membrane. Average fluorescence intensity is quantified along the representative yellow line (drawn from "I" – intracellular – to "E" – extracellular) spanning the plasma membrane.

**Figure S1:** Fluorescence intensity line profiles are taken from at least 5 independent xy-locations in multiple cells, and shaded regions indicate a single standard deviation. Scale bars 10 μm. **B** To account for potential effects on final hPRLR localization due to the signal peptide (SP) sequence, the rat PRLR signal peptide is replaced by the human PRLR signal peptide and the human CD86 signal peptide, mNeonGreen is used as the fluorescent protein (FP) fusion, and the extracellular and intracellular domains of PRLR are fused to the FP. Intracellular hPRLR localization is observed with each signal peptide and configuration of the PRLR-FP fusion. Scale bars 20 μm. Images are maximum intensity projections.

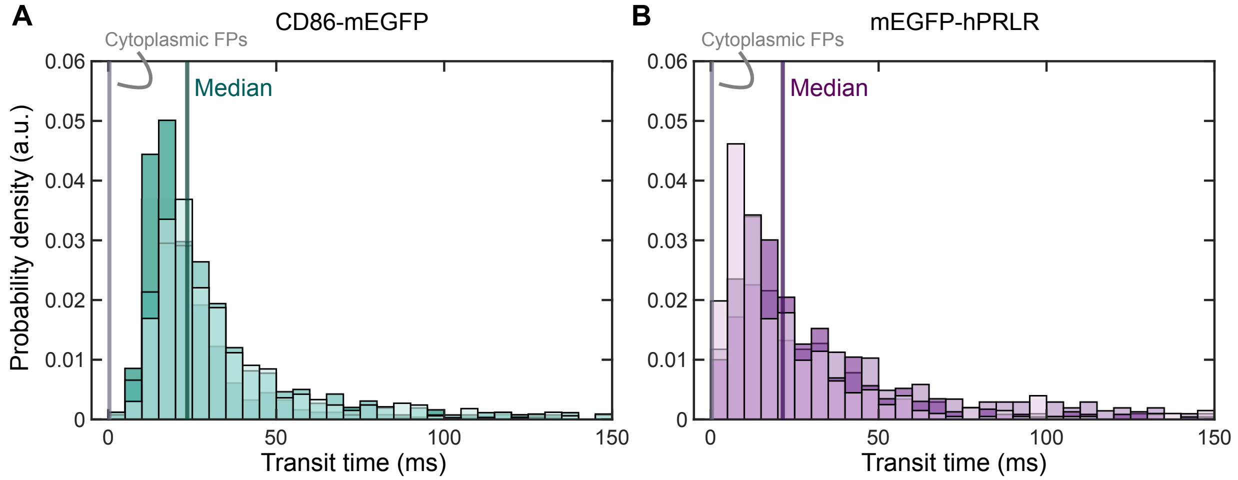


**Figure S2: sFCS replicates. A,B** Transit time histograms for three independent replicates of sFCS measurements taken for CD86-mEGFP and mEGFP-hPRLR. Median transit times across all three replicates are displayed as a vertical line for each protein.

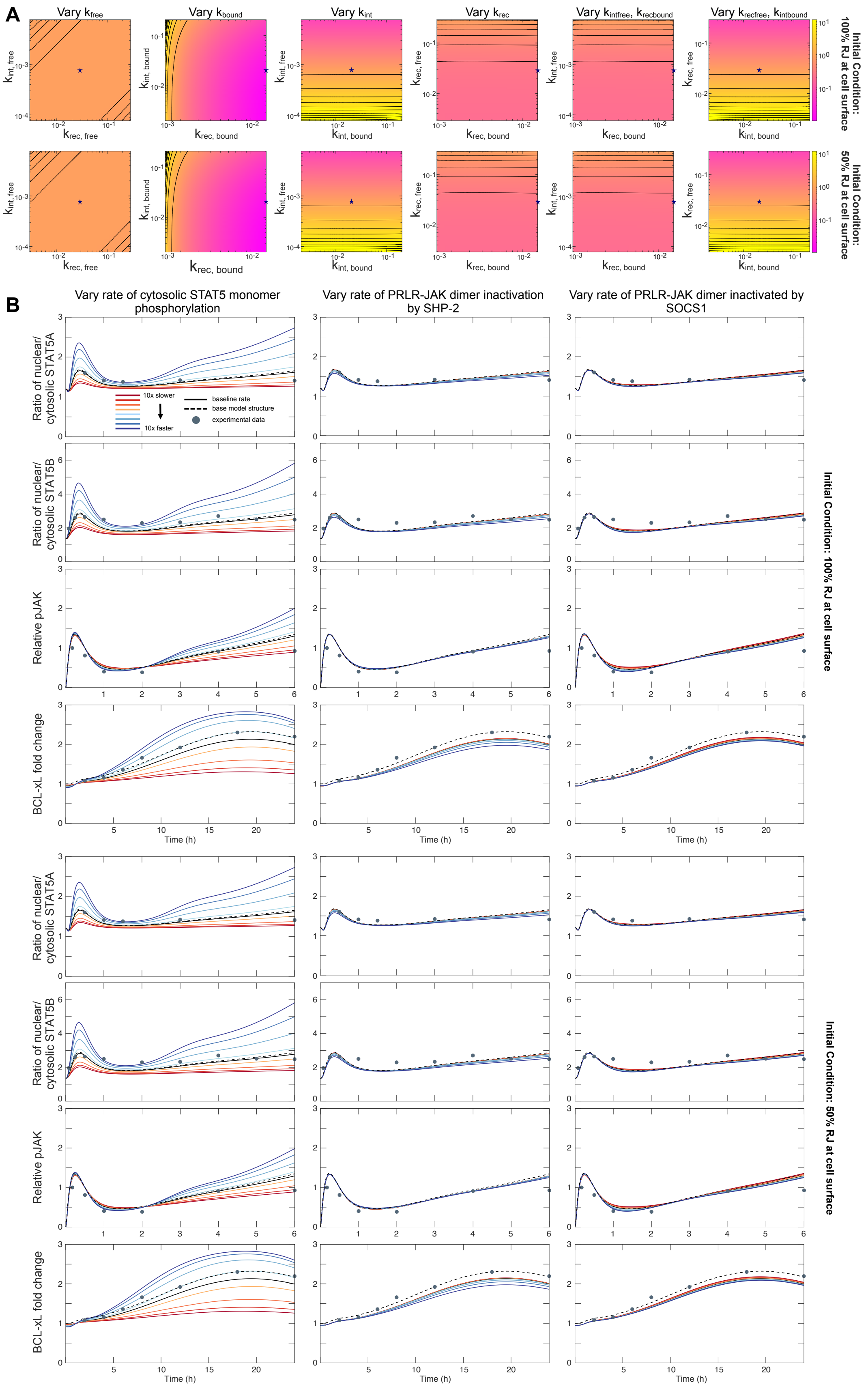


**Figure S3: Model predictions show low sensitivity to the initial spatial distribution of PRLR.**

**Figure S3: A** Heatmaps for meshgrid simulations displaying the receptor localization ratio indicate that initial localization of PRLR does not qualitatively affect the distribution of active receptor complexes after 6h. Representative results are shown for the initial conditions of all RJ starting material localized to the cell surface and 50% of RJ starting material localized to the surface and 50% localized to the intracellular compartment. **B** Tuning downstream signaling rates ($k6_{internal}$, $k10_{internal}$, $k21_{internal}$) while varying the initial distribution of PRLR also shows no qualitative difference over 6h post-prolactin stimulation. Representative results are shown for the initial conditions of all RJ starting material localized to the cell surface and 50% of RJ starting material localized to the surface and 50% localized to the intracellular compartment.

**Table S1. PRLR Trafficking Reaction Network**

**Symbols**

| | |
|---|---|
| $\rightarrow$ | Irreversible reaction |
| $\leftrightarrow$ | Reversible reaction |
| [A:B] | Species complex |
| c | Cytosolic species |
| n | Nuclear species |
| $x_p$ | Phosphorylation |
| i | Internalized receptor species or complex |

| **Reactions** | **Forward Parameter** | **Backward Parameter** |
|---|---|---|
| **Early Receptor Activation** | | |
| $\rightarrow$ RJ | k1 | |
| PRL + RJ $\leftrightarrow$[PRL:RJ] | k2 | k_2 |
| [PRL:RJ] + [PRL:RJ] $\leftrightarrow$ [PRL:RJ2] | k3 | k_3 |
| [PRL:RJ2] $\rightarrow$ [PRL:RJ2a] | k4 | |
| [PRL:RJ2a] + S5c $\leftrightarrow$ [PRL:RJ2a:S5c] | k5 | k_5 |
| [PRL:RJ2a:S5c] $\rightarrow$ $S5c_p$ + [PRL:RJ2a] | k6 | |
| $S5Ac_p$ + $S5Ac_p$ $\leftrightarrow$ [$S5Ac_p$:$S5Ac_p$] | k8A | k_8A |
| $S5Bc_p$ + $S5Bc_p$ $\leftrightarrow$ [$S5Bc_p$:$S5Bc_p$] | k8B | k_8B |
| $S5Ac_p$ + $S5Bc_p$ $\leftrightarrow$ [$S5Ac_p$:$S5Bc_p$] | k8AB | k_8AB |
| **Cytosolic Receptor Inactivation** | | |
| [PRL:RJ2a] + SHP $\leftrightarrow$ [PRL:RJ2a:SHP] | k9 | k_9 |
| [PRL:RJ2a:SHP] $\rightarrow$ [PRL:RJ2] + SHP | k10 | |
| Xpc + [$S5c_p$:$S5c_p$] $\leftrightarrow$ [Xpc:S5c:S5c] | k11 | k_11 |
| Xpc + $S5c_p$ $\leftrightarrow$ [Xpc:S5c] | k11 | k_11 |
| [Xpc:S5c:S5c] $\rightarrow$ Xpc + S5c + S5c | k12 | |
| [Xpc:S5c] $\rightarrow$ Xpc + S5c | k12 | |
| **STAT5 Nuclear Import, Inactivation, and Export** | | |
| $S5c_p$ + S5c $\leftrightarrow$ [$S5c_p$:S5c] | k13 | k_13 |
| [$S5Ac_p$:$S5Ac_p$] $\rightarrow$ [$S5An_p$:$S5An_p$] | k14A | |

| **Reactions** | **Forward Parameter** | **Backward Parameter** |
| --- | --- | --- |
| $[S5Bc_p{:}S5Bc_p] \rightarrow [S5Bn_p{:}S5Bn_p]$ | k14B | |
| $[S5Ac_p{:}S5Bc_p] \rightarrow [S5An_p{:}S5Bn_p]$ | k14AB | |
| $Xpn + [S5n_p{:}S5n_p] \leftrightarrow [Xpn{:}S5n{:}S5n]$ | k15 | k_15 |
| $Xpn + S5n_p \leftrightarrow [Xpn{:}S5n]$ | k15 | k_15 |
| $[Xpn{:}S5n{:}S5n] \rightarrow Xpn + S5n + S5n$ | k16 | |
| $[Xpn{:}S5n] \rightarrow Xpn + S5n$ | k16 | |
| $S5Ac \leftrightarrow S5An$ | k17A | k_17A |
| $S5Bc \leftrightarrow S5Bn$ | k17B | k_17B |
| **mRNA Transcription and Translation; mRNA and Protein Degradation** | | |
| $[S5n_p{:}S5n_p] \rightarrow mRNAn + [S5n_p{:}S5n_p]$ | k18a (Vmax), k18b (Km) | |
| $mSOCS1n \rightarrow mSOCS1c$ | k19 | |
| $mSOCS1c \rightarrow SOCS1 + mSOCS1c$ | k20 | |
| $SOCS1 + [PRL{:}RJ2a] \leftrightarrow [SOCS1{:}PRL{:}RJ2a]$ | k21 | k_21 |
| $mSOCS1 \rightarrow$ | k22 | |
| $SOCS1 \rightarrow$ | k23 | |
| $[SOCS1{:}PRL{:}RJ2a] \rightarrow$ | k24 | |
| $[S5n_p{:}S5n_p] \rightarrow mRn + [S5n_p{:}S5n_p]$ | k25a (Vmax), k25b (Km) | |
| $mRn \rightarrow mRc$ | k26 | |
| $mRc \rightarrow$ | k27 | |
| $mRc \rightarrow Rc + mRc$ | k28 | |
| $Rc \rightarrow RJ$ | k29 | |
| $[S5n_p{:}S5n_p] \rightarrow mBCLn + [S5n_p{:}S5n_p]$ | k30a (Vmax), k30b (Km) | |

| Reactions | Forward Parameter | Backward Parameter |
|---|---|---|
| mBCLn → mBCLc | k31 | |
| mBCLc → | k32 | |
| mBCLc → BCL + mBCLc | k33 | |
| BCL → | k34 | |
| RJ →, iRJ → | kdeg | |
| [PRL:RJ] →, [PRL:RJ2] →, [PRL:RJ2a] → | deg_ratio * kdeg | |
| [PRL:iRJ] →, [PRL:iRJ2] →, [PRL:iRJ2a] → | deg_ratio * kdeg | |
| **Trafficking** | | |
| RJ ↔ iRJ | $k_{int, free}$ | $k_{rec, free}$ |
| [PRL:RJ] ↔ [PRL:iRJ] | $k_{int, bound}$ | $k_{rec, bound}$ |
| [PRL:RJ2] ↔ [PRL:iRJ2] | $k_{int, bound}$ | $k_{rec, bound}$ |
| [PRL:RJ2a] ↔ [PRL:iRJ2a] | $k_{int, bound}$ | $k_{rec, bound}$ |
| [PRL:RJ2a:S5Ac] ↔ [PRL:iRJ2a:S5Ac] | $k_{int, bound}$ | $k_{rec, bound}$ |
| [PRL:RJ2a:S5Bc] ↔ [PRL:iRJ2a:S5Bc] | $k_{int, bound}$ | $k_{rec, bound}$ |
| [PRL:RJ2a:SHP] ↔ [PRL:iRJ2a:SHP] | $k_{int, bound}$ | $k_{rec, bound}$ |
| [SOCS1:PRL:RJ2a] ↔ [SOCS1:PRL:iRJ2a] | $k_{int, bound}$ | $k_{rec, bound}$ |
| [PRL:RJ2a:S5Ac:SHP] ↔ [PRL:iRJ2a:S5Ac:SHP] | $k_{int, bound}$ | $k_{rec, bound}$ |
| [PRL:RJ2a:S5Bc:SHP] ↔ [PRL:iRJ2a:S5Bc:SHP] | $k_{int, bound}$ | $k_{rec, bound}$ |
| [SOCS1:PRL:RJ2a:SHP] ↔ [SOCS1:PRL:iRJ2a:SHP] | $k_{int, bound}$ | $k_{rec, bound}$ |
